# Sequence features do not drive karyotypic evolution: what are the missing correlates of genome evolution?

**DOI:** 10.1101/2022.08.05.502633

**Authors:** Thomas D. Brekke, Alexander S. T. Papadopulos, Martin T. Swain, John F. Mulley

**Author notes:** To whom correspondence should be addressed: Thomas Brekke.

## Abstract

Genome rearrangements are prevalent across the tree of life and even within species. After two decades of research, various suggestions have been proposed to explain which features of the genome are associated with rearrangements and the breakpoints between rearranged regions. These include: recombination rate, GC content, repetitive DNA content, gene density, and markers of chromatin conformation. Here, we use a set of six aligned rodent genomes to identify regions that have not been rearranged and characterize the breakpoint regions where rearrangements have occurred. We found no strong support for any of the expected correlations between breakpoint regions and a variety of genomic features previously identified. These results call into question the utility and repeatability of identifying chromatin characteristics associated with rearranged regions of the genome and suggest that perhaps a different explanation is in order. We analyzed rates of karyotypic evolution in each of the six lineages and found that the Mongolian gerbil genome has had the most rearrangements. That gerbils exhibit very rapid sequence evolution at a number of key DNA repair genes suggests an alternative hypothesis for patterns of genome rearrangement: karyotypic evolution may be driven by variation at a few genes that control the repair pathway used to fix double-stranded DNA breaks. Such variation may explain the heterogeneity in the rates of karyotypic evolution across species. While currently only supported by circumstantial evidence, a systematic survey of this hypothesis is now warranted.

## Introduction

Chromosomal rearrangement is a fundamental evolutionary process leading to karyotypic variation that can be found at every taxonomic level from large-scale differences between divergent taxa (Takagi and Sasaki 1974; Graphodatsky et al. 2011; Ferguson-Smith and Trifonov 2007) to chromosomal races within a single species (Piálek et al. 2005). Chromosomal rearrangements facilitate adaptation by locking together coadapted gene complexes (Joron et al. 2011), drive speciation by suppressing fertility when heterozygous (Coyne et al. 1993; Rieseberg 2001), and are implicated in a variety of disease states and cancers (Bailey 2006; Mitelman 2000). As our ability to sequence and assemble entire chromosomes matures and the quality of sequenced genomes increases, the field of genomics has been searching for reasons behind genomic rearrangements. What causes a chromosome to break at a particular site? Are chromosomes prone to repeated breaks at the same site, and what might characterize these regions? It remains unknown whether rearrangements are driven by specific features (e.g., unusual GC content or repetitive elements), epigenetic marks and chromatin conformations, or simply biases in the repair of double strand breaks.

Advances in understanding the evolution of rearrangements have closely followed the development of technical and methodological techniques. Studies from the 1980s that use comparative mapping approaches were only able to look at the size distribution of unbroken regions (Nadeau and Taylor 1984) while cytogenetic staining of chromosomes could co-localize breaks with banding patterns (Glover and Stein 1988; Sutherland et al. 1985). As full-genome sequencing came online in the early 2000s, the focus shifted to testing for correlations between primary sequence features such as GC content or gene density with breakpoint regions (for an in-depth review see (Farré et al. 2015)). By the late 2000s a sufficient number of genomes were assembled to chromosome scales that the role of recombination on rearrangements could be investigated (Larkin et al. 2009; Farré et al. 2013; Ullastres et al. 2014; Capilla et al. 2016). Methods were developed in the 2010s that could analyze the conformation, nuclear location, and epigenetic state of chromosomes and these led to a slew of papers linking rearrangements with higher level structural and regulatory features (reviewed in (Farré et al. 2015)). Throughout this time positive correlations were identified at all levels: from GC content of rearranged regions to chromatin conformation, and from recombination hotspots to epigenetic modifications. To synthesize this breadth of phenomena the ‘Integrative Breakage Model’ (IBM) was proposed which attempts to integrate effects of the physical characteristics of the genome including chromatin position in the nucleus, chromatin structure, and epigenetic variation with the selective constraints of avoiding disruption to functional genes and regulatory blocks (Farré et al. 2015).

While a unified theory encapsulating nearly 40 years of research is a necessary advance, the findings that underlie the formation of the IBM are not as unified as they first appear. The only clear consensus to date is that recombination rate is typically lower inside breakpoint regions than outside (Larkin et al. 2009; Farré et al. 2013; Ullastres et al. 2014; Capilla et al. 2016), otherwise evidence for the genomic correlates of breakpoints is, at best, scattered and contradictory (Table 1, Table S1). Various types of sequences are often enriched in breakpoint regions, but different studies find enrichment for different sequence motifs: some find enrichment for centromeric sequences (Zhao et al. 2004; Mlynarski et al. 2009) or for various transposons (Thybert et al. 2018; Penso-Dolfin et al. 2020). Some find enrichment for telomeric sequences (Eichler and Sankoff 2003; Paço et al. 2013) and others find no enrichment for telomeric sequences (Murphy et al. 2005). As per the IBM, open chromatin seems important but the studies underlying this conclusion all measured different features. The model therefore rests on the assumption that open chromatin state correlates with distance to the origin of replication (Lemaitre et al. 2009), transcription factor binding sites (Farré et al. 2019), CpG islands (Larkin et al. 2009), hypomethylaion of CpG islands (Carbone et al. 2009; Lemaitre et al. 2009), transcription level (Lemaitre et al. 2009; Capilla et al. 2016), lamina-associated domains, active chromatin histone marks, CTCF sites, RNA pol II sites, and DNaseI hypersensitivity sites (Capilla et al. 2016). While this evidence from chromatin state all points in generally the same direction, results concerning the parts of the genome that may constrain evolution through negative selection (e.g.: genes, regulatory regions, etc) are contradictory. We would expect that regions of the genome under strong purifying selection would be unlikely to act as rearrangement hotspots and yet some studies have reported high gene density in breakpoint regions (Murphy et al. 2005; Ma et al. 2006; Kemkemer et al. 2009; Larkin et al. 2009; Lemaitre et al. 2009; Ullastres et al. 2014; Capilla et al. 2016; Farré et al. 2016). Others report low gene density in breakpoints (Kikuta et al. 2007; Becker and Lenhard 2007) or that it depends on the regulatory complexity of the genes themselves (Mongin et al. 2009). Topologically associated domains (TADs) also present a bit of a puzzle. We would expect that these large-scale regulatory blocks would resist a rearrangement within them, and indeed it is the boundaries in between the TADs that are closely associated with breaks in synteny when comparing humans and gibbons (Lazar et al. 2018). This pattern would be expected if disrupting TADs alters gene expression and is selected against. However, recent experimental evidence has come out that breaking TADs does not actually change gene expression, at least in *Dropsophila* (Ghavi-Helm et al. 2019) and so it is not clear why rearrangements seemed to avoid breaking the TADs of the human-gibbon ancestor.

**Table 1:** Summary of studies looking at genomic correlates of rearrangements. * “Things that may cause evolutionary constraing in BPs vs genome” include conserved non-coding elements, genes, and TADs, further details of all of these categories can be found in Table S1.

| Citation | Year | Recombination rate in BPs vs genome | Density of sequence motifs in BPs vs genome | Chromatin conformation and epigenetics in BPs | Density of things that may cause evolutionary constraints in BPs vs genome |
| --- | --- | --- | --- | --- | --- |
| Eichler and Sankoff | 2003 | - | Various motifs high | - | - |
| Zhao et al. | 2004 | - | Centromeres high | - | - |
| Murphy, et al. | 2005 | - | Various motifs high, telomeres low | - | High |
| Weber and Ponting | 2005 | - | GC content high | - | - |
| Ruiz-Herrera | 2006 | - | Segmental duplications high | - | - |
| Ma, et al. | 2006 | - | Various motifs high | - | High |
| Kikuta | 2007 | - | - | - | Low |
| Becker | 2007 | - | - | - | Low |
| Kemkemer et al. | 2009 | - | Segmental duplications high | - | High |
| Mongin | 2009 | - | - | - | Mixed |
| Larkin, et al. | 2009 | Low | Segmental duplications high | Open | High |
| Lemaitre et al. | 2009 | - | GC content high | Open | Mixed |
| Carbone, et al. | 2009 | - | Various motifs high | Open | - |
| Mlynarski, et al. | 2010 | - | Centromeres high | - | - |
| Farre et al | 2013 | Low | - | - | - |
| Paco et al | 2013 | - | Telomeres high | - | - |
| Ullastres, et al | 2014 | Low | - | - | High |
| Capilla, et al. | 2016 | - | - | Open | High |
| Farre et al. | 2016 | - | - | - | Mixed |
| Damas, et al. | 2018 | - | - | - | Low |
| Thybert, et al. | 2018 | - | Transposons high | - | - |
| O'Connor, et al. | 2018 | - | - | - | Low |
| Farre, et al. | 2019 | - | Transposons high, no enrichment for segmental duplications | Open | - |
| Penso-Dolfin, et. al. | 2020 | - | Transposons high | - | - |
| this paper | 2020 | Low | No difference | No difference | No difference |

The fundamental issue apparent in the history of the field is the twofold failure to replicate and to report negative findings. This combination of issues makes interpretation of any overarching pattern difficult. Indeed, out of the 26 studies, any given genomic feature was discussed on average in 3.4 papers (median = 2). Thus, nearly every genomic feature was found to be important by a few studies but was not considered by most. Irregular taxonomic sampling has further complicated the issue: are segmental duplications really the driving factor for genome rearrangements in birds and mammals (Larkin et al. 2009) except for ruminants (Farré et al. 2019)? Do CpG islands only influence rearrangements in primates (Larkin et al. 2009)? And is gene content key because gene density is high in mammals (Ma et al. 2006; Murphy et al. 2005; Kemkemer et al. 2009; Larkin et al. 2009; Lemaitre et al. 2009), but because gene density is low in fish (Kikuta et al. 2007; Becker and Lenhard 2007)? The forces that govern karyotypic evolution appear to vary drastically by lineage, but there is no theoretical basis to explain these inconsistencies.

Much of the disparity in previous results stems from the assumption that whatever is unique about a genome is so because it was selected on. This fallacy can easily lead to a variety of conflicting results as we describe above. The synthesis papers that have attempted to find a general model for genome evolution have taken all positive results as reported and then been left with the conclusion that the forces that govern genome evolution are hugely varied and lineage-specific (Farré et al. 2015). Here we evaluate the methods of previous studies critically by replicating the approach on a similarly sized dataset. We use a hypothesis-driven approach based on first principles derived from current theory in molecular biology, genetics, and evolution. We formulate and test hypotheses about the roles that (1) recombination, (2) repetitive sequence motifs, (3) evolutionary constraints on genes and regulatory blocks, and (4) chromatin conformation play in genome rearrangements. To test our predictions, we use full genome alignments between six rodents (house mouse - *Mus musculus*, rat - *Rattus norvegicus*, Mongolian gerbil - *Meriones unguiculatus*, deer mouse - *Peromyscus maniculatus*, meadow vole - *Microtus orchogaster*, and Chinese hamster *-Cricetulus griseus)* from which we identified evolutionary breakpoint regions. We failed to find any results consistent with previous studies. Consequently, it is likely that searching for the sequence-level correlates of genome rearrangements using widely applied approaches is a problematic and we offer a possible alternative hypothesis to explain the distribution of breakpoints.

## Results and Discussion

### Syntenic Fragments and breakpoint regions

Using full-genome alignments between six rodent species, we identified 317 syntenic fragments (SFs; Fig S1 and Dataset S1). These fragments range in size from 200Kb to 80Mb and nearly always contain multiple, and often numerous, topologically associating domains (TADs) which are typically less than 100Kb long (Liu et al. 2019). The length of the breakpoint regions (BPs) between these fragments vary from 1 base to 11Mb (Fig S2). Both the location and identity of our breakpoints generally agree with those described by Mlynarski (Mlynarski et al. 2009) which were based on chromosome banding patterns.

We used these syntenic fragments to test whether a variety of genome features are correlated with genomic breaks in two complimentary ways (Fig S3). First, using house mouse, rat, deer mouse, and vole we compared features between syntenic fragments and the breakpoint regions. Chinese hamster and Mongolian gerbil were omitted from these types of analyses as they lack chromosome-scale genomes and therefore the sequences of the breakpoint regions are unknown. Second, we compared the incidence of the genomic feature across each SF under the assumption that there should be some enrichment (or de-enrichment) near the end of the fragment where the breakpoint occured; a strategy with less power, but one that works in all six species because the sequence of the breakpoint regions is not required. The results of these analyses for each genomic feature are discussed below.

### Recombination and rearrangements

A double stranded DNA break must initiate any novel genome rearrangement. This observation leads to the hypothesis that rearranged sites may ultimately derive from a mistake during crossing-over in meiosis. Therefore, we predicted that rearrangements occur more often in regions of the genome with high recombination rates. To test whether the recombination rate was higher in breakpoint regions than syntenic fragments of the four species with chromosome-scale genomes, we used a linear mixed effects model with species as the random effect. Counter to our hypothesis, the best model shows that the recombination rate is consistently lower in breakpoint regions than in syntenic fragments across species (Fig 1A, χ^2^ _df=1_ = 87.091, p < 0.00001), though the effect size of breakpoint region-vs-syntenic fragment (1.78 cM/Mb) is smaller than the residual variation (2.38 cM/Mb). To test whether the recombination rate is higher close to breakpoint regions we examined recombination rate in 1kb windows along 10Mb sections of syntenic fragments adjacent to the breakpoint regions. There is a great deal of variation across species: *Meriones* and *Microtus* have a negative relationship between recombination rate and distance from a breakpoint while *Mus, Rattus*, and *Peromyscus* all have positive relationships (Fig S4). However, a mixed effects model with species as a random effect and distance to the breakpoint as the fixed effect showed a significant positive relationship and outperformed the null model of species as a random effect (χ^2^ _df=1_ = 2159.3, p < 0.00001). These results - lower recombination rates in and around breakpoint regions - are in line with previous findings in rodents (Larkin: 2009ij; Capilla et al. 2016). Recombination occurs in hotspots in many mammal genomes (Steinmetz et al. 1987) and that the location of these hotspots can change rapidly through evolutionary time (Coop and Myers 2007), though modelling suggests that defunct hotspots may occasionally be resurrected (Úbeda et al. 2019). If rearrangements occur at evolutionarily labile recombination hotspots, then the relationship between the present recombination rate and past rearrangements may be weak (Farré et al. 2015), which may go some way towards explaining the observed pattern.

**Figure 1:**
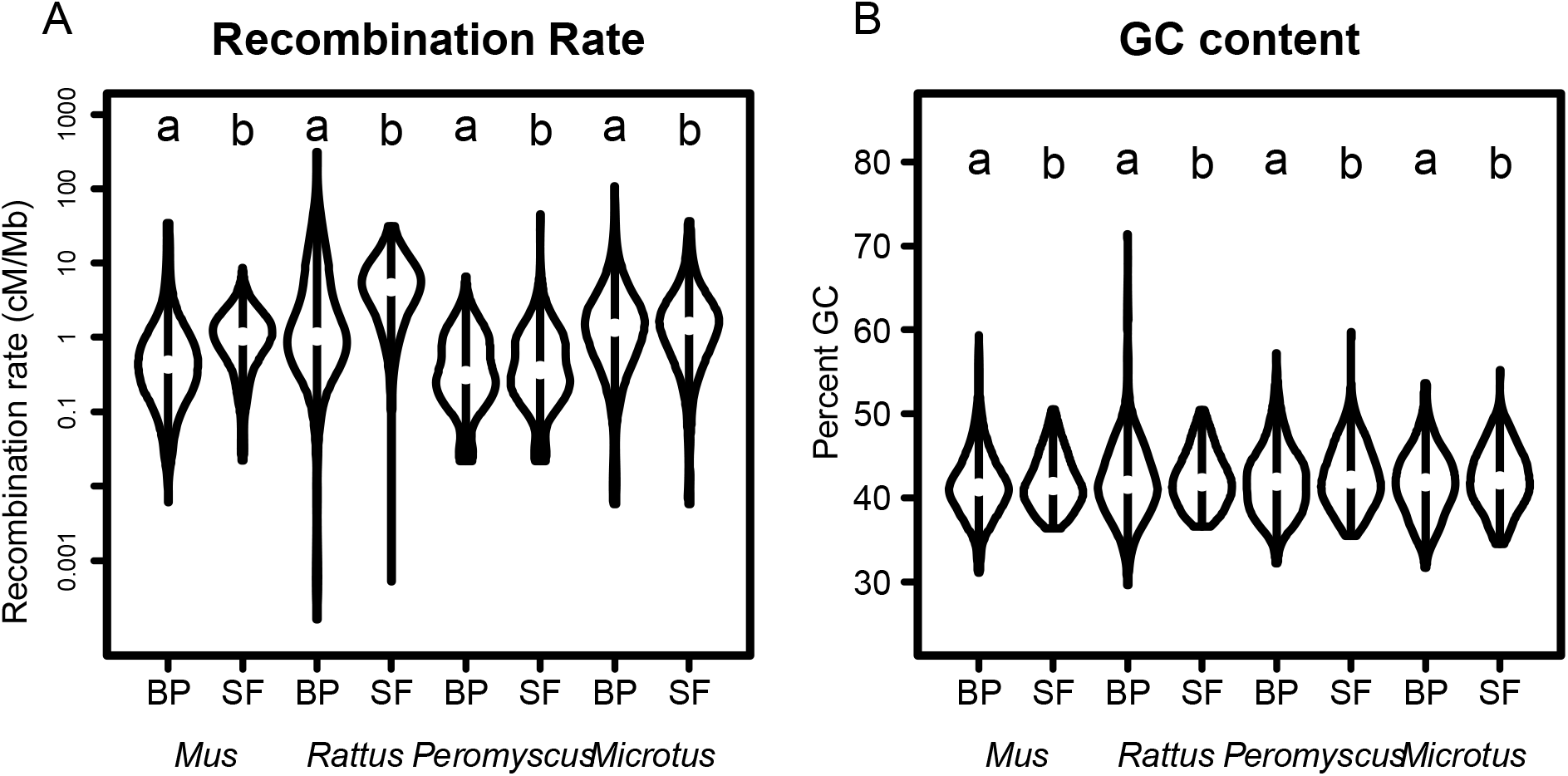
Recombination rate and GC content differences between syntenic fragments and breakpoint regions. The white point marks the median value and the thick black bar marks the interquartile range. (A) Recombination rate is lower in breakpoint regions (BP, grouping ‘a’) than in syntenic fragments (SF, grouping ‘b’, χ^2^ _df=1_ = 87.091, p < 0.00001). If meiotic double-strand breaks drive rearrangements we expect to see just the opposite pattern. Similarly, recombination rate within SFs tends to increase with distance from a breakpoint (Fig. S4, χ^2^ _df=1_ = 2159.3, p < 0.00001) though there is much species-level variation. (B) GC content is slightly, but consistently 0.36% lower in breakpoint regions as well (χ^2^ _df=1_ = 5.5681, p = 0.01829). These data too, are counter to the prediction that recombination rate and hence GC content should be higher in rearranged areas.

Though the recombination rate may vary, elevated GC content (a feature commonly associated with recombination hotspots), should be more stable through evolutionary time. GC content rapidly increases near recombination hotspots due to biased gene conversion favouring Gs and Cs during the mismatch repair pathway (Arbeithuber et al. 2015). But once the GC content is high and the hotspot moves or dies there is little evolutionary pressure on GC content directly and the GC levels may remain elevated through evolutionary time. Therefore, a correlation between breakpoint regions and GC content may be a more reliable indicator of recombination-mediated rearrangement. This pattern has been reported previously (Ma et al. 2006; Webber and Ponting 2005), but we found no evidence for it here: our breakpoint regions contain 0.36±0.15(se) % fewer GC sites than syntenic fragments (Fig 1B, χ^2^ _df=1_ = 5.5681, p = 0.01829). The effect size is small but is consistent across all species. Similarly, when we regress GC content across the 100kb next to breakpoint regions, there is no overall trend, only some small differences between species (Fig S5, χ^2^_df=1_ = 0.377, p = 0.846). Taken together, the recombination rate and GC content data indicate that genomic breakpoints are unlikely to be the result of mistakes during crossing over because otherwise the locations of the rearrangements would correlate with recombination rate. Rather, it suggests that rearrangements are instead due to incidental DNA damage which is repaired incorrectly and differences in rates of karyotypic evolution may therefore be the result of variation in the fidelity of the DNA repair pathways in different species.

### Sequence motifs and rearrangements

Features of the DNA sequence such as repetitive elements may be enriched in breakpoint regions for one of two complementary reasons: 1) repetitive sequence motifs may be inherently fragile and predispose specific areas to breakages and 2) once a chromosome has broken, similar repeats in different areas of the genome may provide the micro-homology with which to resolve the double-strand break (Villarreal et al. 2012; Wang and Xu 2017). Indeed, sequence motifs like those in telomeres (Paço et al. 2013), centromeres (Murphy et al. 2005; Mlynarski et al. 2009), or transposons (Ma et al. 2006; Carbone et al. 2009; Farré et al. 2019; Penso-Dolfin et al. 2020) may be enriched in rearrangement breakpoint regions. Our goal was to test whether repetitive sequences are found in breakpoint regions more often than not and to do so using a test that would be robust across a variety of repeat types and lengths. If there is some sequence correlated with rearrangements, then we would expect that the kmer spectra distances between breakpoint regions would be low overall (i.e., breakpoint regions should have similar sequence composition to other breakpoint regions, whereas, SFs should not necessarily be similar to other SFs). We used a linear mixed effects model to test for the effect of comparison type (BP vs BP, BP vs SF, or SF vs SF) on the distance with species, kmer length, and method as random effects. This model was significantly better at explaining the data than a null model with only the random effects (Figure 2 and Fig S6, χ^2^ _df=2_ = 928, p < 0.00001) and it showed that the least variation (i.e., lowest distance) is found when comparing breakpoint regions. This is the expected pattern if there is some common sequence motif present in breakpoint regions. However, the residual variance in the model (0.273) is much larger than the effect size of either BP-SF (0.067) or SF-SF (0.055) suggesting that the ostensible similarity of breakpoint regions is slight. While statistically significant, these data do not convincingly demonstrate the presence of consistent repetitive sequence which correlates with breakages.

**Figure 2:**
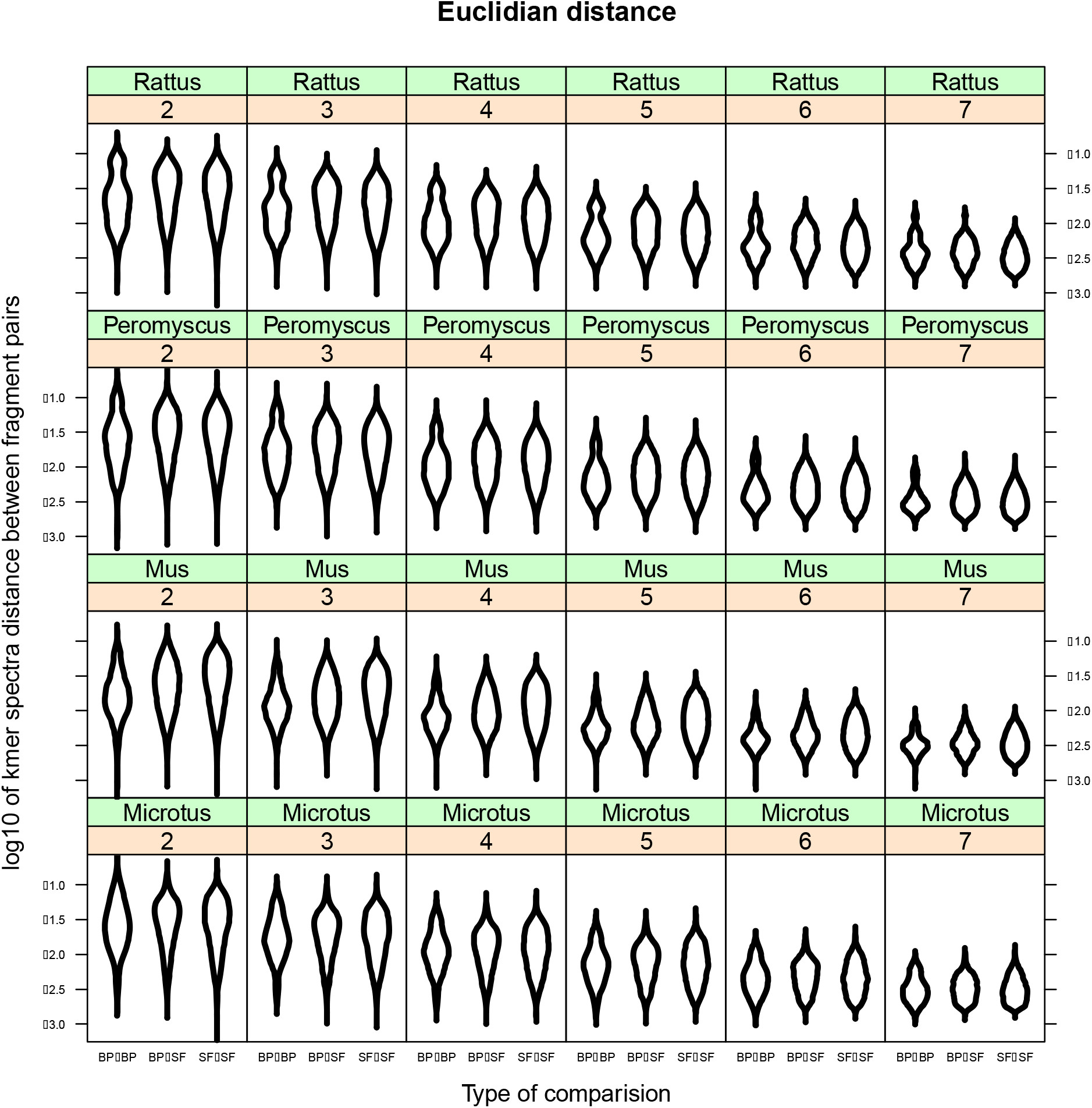
Kmer analysis and sequence similarity across breakpoint regions. Kmer spectra differences measured with Euclidian distance using kmers of size 2, 3, 4, 5, 6, and 7 (number in orange boxes). The lowest variation is found when breakpoint regions are compared with themselves (‘BP-BP’) while the variation with syntenic fragments (‘BP-SF’ and ‘SF-SF’) is higher (χ^2^_df=2_ = 928, p < 0.00001). This figure shows the Euclidean distance metric and the same pattern is found in the other two methods used to estimate variation as well (d2 and Chebyshev) (Fig. S5). These data are consistent with the hypothesis that breakpoint regions share some sequence similarity (i.e. telomere repeats, centromere repeats, microsatellites, transposons, etc). However, the residual variation (0.273) is far greater than the effect size (‘BP-SF’ effect size = 0.067, ‘SF-SF’ effect size = 0.055) suggesting the presence of some other important factor which is not accounted for.

### Evolutionary constraints on rearrangements

Distinct from the question of whether specific features of genome attract or facilitate breaks, are questions about which sorts of rearrangements can become fixed in a population. Novel arrangements start as single copies and research suggests that they are often underdominant with a potentially serious selective cost (Stathos and Fishman 2014). Early research suggested that they can fix by drift in a very small populations (Rieseberg 2001) or by positive selection if they tie together locally adapted alleles (Kirkpatrick and Barton 2006). By linking multiple locally adapted alleles, the positive selection that results from keeping those alleles together (i.e., reduced recombination) can overcome the selective disadvantage of the rearrangements’ underdominance. Such local adaptation can drive speciation and is aided by both large effective population size and by population substructure (Rieseberg 2001). Indeed, mammalian taxa with large litter sizes (which are typically also those with large population sizes) have higher rates of karyotype evolution (Martinez et al. 2017) implying that positive selection may drive the fixation of novel rearrangements. In addition to locking together locally adapted allele complexes, rearranged areas may experience positive selection as novel gene function arises from breakup and reassembly of old genes (Larkin et al. 2009).

Any mutational process that disrupts functional genes should generally be selected against. If rearrangements are generally deleterious we predict that: (i) the breaks that have fixed should be found in relatively gene-poor regions where they did not disturb the function, and (ii) the genes that do occur in breakpoint regions should be short as they would be more likely to survive the rearrangement process intact. Similarly, and at a slightly larger scale, we would not expect to find breakpoints in the middle of co-regulated gene complexes such as TADs. Alternatively, a high density of long genes in breakpoint regions may suggest that either positive selection on novel exon arrangements is a predominant factor driving the fixation of novel rearrangements or that chromosome breaks are more likely to occur in regions of the genome with open chromatin being actively transcribed.

To test these predictions, we have analysed both gene density and gene length in breakpoint regions compared with syntenic fragments. We used two different metrics of gene density - either counting all bases between the beginning of the 5’ UTR and the end of the 3’ UTR or counting only exonic bases. For each of these tests we used a linear mixed effects model with species as a random effect. We find that breakpoint regions are not more enriched for genic bases (including UTRs and introns) than syntenic fragments (Figure 3A, χ^2^ _df=1_ = 1.36, p = 0.2512). We also regressed gene density across the syntenic fragment using a similar linear mixed effects model to confirm the pattern. Surprisingly, the model including distance to the breakpoint region was significantly better than an intercept-only model (Fig. S7A, χ^2^ _df=1_ = 11103, p < 0.00001) and has a negative trend of genes per base moving away from breakpoint regions. This implies that breakpoint regions may have higher gene density than syntenic fragments in contrast to our simpler test above. However, the effect size is vanishingly slight: a decrease of 0.003 genic bases per megabase. These results suggest that genome-wide, the balance of selection against deleterious rearrangements and for advantageous arrangements is broadly similar to that of other new mutations.

**Figure 3:**
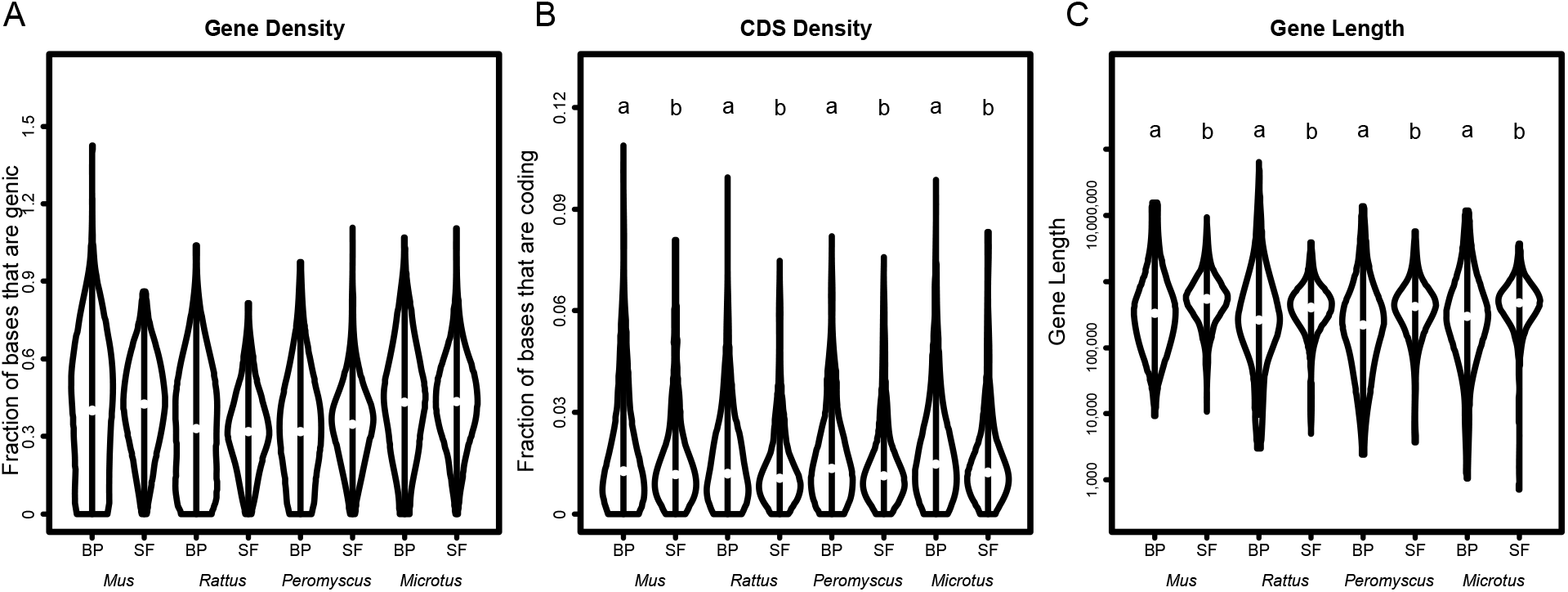
Gene density and length. (A) There is no difference in the fraction of genic bases in syntenic fragments and breakpoint regions (χ^2^ _df=1_ = 1.36, p = 0.2512). (B) However, when measuring only coding bases, there is a significant trend where breakpoint regions have a higher proportion of CDS bases than syntenic fragments (χ^2^ _df=1_ = 9.849, p = 0.001699). The effect size is slight though (1.87 more CDS bases per kb) especially when compared to the variation across samples (14.8 bases per kb). Furthermore, the opposite pattern is found when regressing gene content across a syntenic fragment (Fig S7). (C) The genes in breakpoint regions are significantly shorter than those in syntenic fragments: breakpoint genes are 25,877 bases long from the beginning of the 5’UTR to the end of the 3’UTR, while genes in syntenic fragments are 42,450 bases long (χ^2^ _df=1_ = 78.323, p < 0.00001).

Selective constraints arise from function and so we also repeated the same pair of analyses using only coding bases. Here a linear mixed effects model with fragment type as the fixed effect and species as a random effect outperformed a simpler model with only species as a random effect (Figure 3B, χ^2^ _df=1_ = 9.849, p = 0.001699). Here too the better model suggests that breakpoint regions have a higher density of coding though the effect size is again quite small compared to the residual variation: breakpoint regions have 1.87 more coding bases per thousand bases. Compared to the residual variation of 14.8 coding bases per thousand bases, a difference of less than 2 coding bases per kb does not seem not biologically meaningful. As breakpoint regions are on average 100kb long, this translates to approximately 200 bases, typically less than one gene, difference between a breakpoint region and a similar-sized piece of an SF. A regression of genic bases across the syntenic fragments tells the same story: a significant decrease in gene density moving away from breakpoint regions. Specifically, a model including distance to the breakpoint outcompeted the null intercept-only model (Fig. S7B, χ^2^ _df=1_ = 4417570, p < 0.00001). The effect size of the superior model is again slight: a decrease of 0.0004 genic bases per megabase. These four tests, evaluating different variations on the same fundamental genome feature, resulted in a mix of outcomes: gene density is either the same or higher in breakpoint regions when compared with syntenic fragments directly and tends to decrease as we move away from breakpoint regions within a syntenic fragment. These results then are in line with some of the previous studies which also found higher gene content in breakpoint regions (Murphy et al. 2005; Larkin et al. 2009; Capilla et al. 2016; Farré et al. 2016) and counter to other studies which found lower gene density (Kikuta et al. 2007; Becker and Lenhard 2007; Damas et al. 2018; O’Connor et al. 2018), but in general the effect sizes we found are so slight we are hesitant to argue either way.

While the gene-density question remains open, breakpoint regions contain genes which tend to be shorter than average. The median gene length (including introns) within breakpoint regions is 25,877 bases, while the median length of those in syntenic fragments is 42,450 bases (Figure 3C, χ^2^ _df=1_ = 78.323, p < 0.00001). These data suggest that the rearrangements that split up the exons of multi-exonic genes do incur a selective cost. Taken together, these two lines of evidence are most consistent with the hypothesis that preserving gene function is an important selective filter that can keep rearrangements from fixing. This interpretation does not preclude the generation of novel genes in breakpoint regions but instead suggests that positive selection on novel gene function is not the primary driving force behind the fixation of rearrangements. In addition, genomic regulatory blocks such as TADs mean that functional genomic units extend far beyond simply the coding bases and likely further constrain the location of the breakpoints that are allowed to fix in the population.

### Chromatin conformation and rearrangements

As rearrangements are fundamentally the result of DNA damage being repaired incorrectly, it is reasonable to expect that chromatin conformations that expose DNA to damage would be enriched in breakpoint regions. Indeed, open chromatin, like that being actively replicated or transcribed, is far more likely to break than tightly-packed inactive chromatin (Fungtammasan et al. 2012). In addition, many studies have found a wide variety of genomic features that are positively correlated with open chromatin enriched in breakpoint regions. These include things related to gene activity such as transcription factor binding sites (Farré et al. 2019), and overall transcription level (Lemaitre et al. 2009; Capilla et al. 2016), the presence of CpG islands (Larkin et al. 2009) and the methylation state of CpG islands (Carbone et al. 2009), active chromatin histone marks, RNA pol II sites, DNaseI hypersensitivity sites, and CTCF sites (Capilla et al. 2016). There are also correlations between replication-related features such as the distance to the origin of replication (Lemaitre et al. 2009), and an inverse correlation between lamina-associated domains (which are inactive chromatin sections) and breakpoint regions (Capilla et al. 2016). All of these features covary with each other across the genome and also with gene density which makes it difficult to distinguish whether chromatin state facilitates breaks (i.e., open DNA that is being bound by transcription factors etc is more likely to break), or whether breaks are only tolerated in certain regions as discussed above. To further complicate the issue: evaluating chromatin state directly in extant animals is expensive and non-trivial; evaluating chromatin state of the extinct ancestor in whose genome the rearrangement occurred is likely impossible. As a proxy for chromatin state, we have tested whether the density of CTCF binding sites vary between BPs and SFs. CTCF binding protein is a major regulator of chromatin architecture which serves as a gene expression insulator and marks boundaries between active and condensed chromatin (Kim et al. 2019). As such we expect that CTCF binding sites should correlate with the boundaries between syntenic fragments and thus be enriched in breakpoint regions. CTCF sites are not well characterised in many species and so we used only house mouse for this test. We find no significant differences between CTCF site density in and out of breakpoint regions in house mouse (Fig. S8, t-test, t_df=642_ = −1.118, p = 0.2639). Thus, we found no evidence for a broad role of chromatin state in the origin and maintenance of rearrangements, but acknowledge that this approach has substantial limitations.

### Genome rearrangements in rodents

Having found no strong candidates for specific genome features that correlate with rearrangements, we next characterise the trajectories of karyotypic evolution in the six focal species as any pattern in the number and type of rearrangements may provide a clue as to the forces governing rearrangements. Some patterns are intuitive: Chinese hamsters have 10 autosomes whereas the other rodents in our study have around 20. Thus, at some point in the evolutionary history of Chinese hamsters a bias has arisen towards chromosomes fusions over chromosome fissions. In contrast, the vole genome is reported to have a high number of inversions (McGraw et al. 2011; Romanenko et al. 2018), suggesting that they are prone to a wholly different mechanism of karyotypic evolution than the hamsters. In addition, house mice are known to be susceptible to Robertsonian translocations across their home-range in Europe (Garagna et al. 2014). Finally, species in the Gerbillinae subfamily are thought to have an increased frequency of rearrangements *(Meriones* (Benazzou et al. 1982), *Taterillus* (Dobigny et al. 2002), and *Gerbillus* (Aniskin et al. 2006)). While these patterns are striking, they are not directly comparable: for instance it is impossible to tell from the above whether gerbils or hamsters experience more rearrangements. To directly compare the pattern of karyotypic evolution in the focal species, we assembled a phylogenetic tree based on the number of rearrangements that separate each genome using the program MGR (Bourque et al. 2004). From the complete set of 317 syntenic fragments, we extracted 115 that were associated with a chromosome and location in all six of our target genomes. In the absence of an *a priori* phylogenetic tree, MGR identified *Microtus* and *Meriones* as the most distinct genomes, clustering together and each with far more changes than any other tip-branch (Figure 4A). When MGR was run in the phylogenetically informed mode the results retained the correct relationship between five of the species including the murids *Mus* and *Rattus* and the three cricetids: *Cricetulus, Peromyscus*, and *Microtus* (Figure 4B). Our data reaffirms previous evidence that vole genomes have many rearrangements (McGraw et al. 2011; Romanenko et al. 2018). Even despite the high number of rearrangements, voles are found in the expected place in the tree: sister to *Cricetulus* with *Peromyscus* as the cricetid outgroup. *Meriones* however, is not placed among the murids as it should be but instead falls out among the cricetids and even so has the most rearrangements. This suggests that the gerbil genome has undergone many rearrangements compared to other murids. Gerbils are unusual regardless of whether the phylogeny informs MGR or not, suggesting that among rodents, gerbil genomes are particularly unstable.

**Figure 4:**
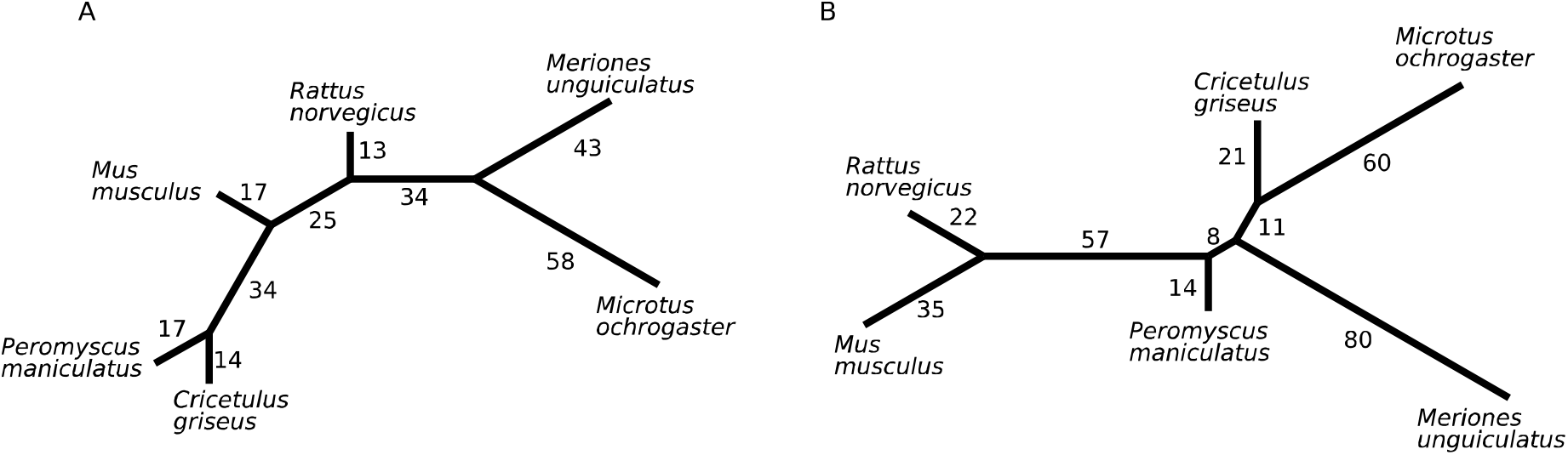
Rearrangement distances between genomes. (A) Phylogeny-naïve unrooted tree. This unrooted phylogeny was built by MGR by taking an ordered set of fragments from a pair of genomes and rearranging them step-by-step until the order of the fragments in each genome are the same. The branches are labeled with the number of rearrangements it takes to transform the order of the fragments at one node to the order at another. The vole genome is known to have many rearrangements presumably drawing it well away from the other cricetids. That the *Meriones* branch is sister to, and nearly as long as the vole branch demonstrates the rampant rearranging in gerbil genomes. (B) MGR was seeded with the known phylogeny of these six species. In this case it properly recovers the relationship between mouse, rat, deer mouse, vole, and hamster but misplaces gerbil among the cricetids. This is likely due to the very high number of rearrangements in the gerbil genome.

The high variation in the number and types of rearrangements across lineages demonstrate that lineage-specific processes must be occurring. However, the argument that different sequence characteristics drive variation in karyotypic evolution requires us to postulate a vast amount of sequence evolution which predisposes some species to chromosome breaks at, for example, regions with centromeric sequences (i.e.: Mynarskyi et al, 2010) while predisposing the breaks in other species to be near, for example, hypomethylated sites (i.e.: Carbone et al 2009), etc. This extent of genome-wide sequence evolution would leave signatures of selection in every genome, but these are conspicuously absent. As a result, the question remains: what sort of evolutionary change could underlie such diversity?

### Reconciliation

A great deal of research has gone into finding genomic correlates of rearranged regions over the last two decades (Table 1) but very few general patterns have been discovered. We argue that the lack of a robust correlation between any given genomic feature and rearrangements is due to the fact that there is no single genome feature that drives rearrangements. The rearrangements we observe between species are the subset of randomly occurring double stranded breaks which preserve gene function and are therefore able survive a selective filter. We suggest that the difference in rates of karyotypic evolution between species would therefore not be due to expansions of a repeat or some other feature that causes rearrangements, but instead to the cellular machinery that repairs DNA double strand breaks. A small amount of evolutionary change in the enzymes responsible for genome maintenance could easily be sufficient to send species down different paths of karyotypic evolution. Indeed a specific instance of this mechanism has been proposed before in relation to novel transposable elements that insert near DNA repair genes in gibbons (Okhovat et al. 2020) and alter the rearrangement landscape by influencing expression of those genes. In the more general case, species and taxa which have highly effective DSB repair pathways should exhibit low rates of karyotypic evolution as the breaks formed are properly repaired, while species that have accumulated mutations (be it transposable elements or otherwise) that inhibit or alter the repair pathway would have more rearrangements, or at least a different rearrangement profile. Furthermore, there are several different DSB repair pathways each of which has its own biases. Mutations in one pathway may change both the rate of karyotypic evolution and the types of rearrangements seen across different species. For instance, one of the major DSB repair pathways is homologous recombination which uses the sister chromosome to accurately repair breaks (Jackson 2002). When the gene *Brca2*, which is central to the homologous recombination repair pathway, is knocked out in mice they accrue many rearrangements as repair is carried out by an alternative pathway: non-homologous end joining (Yu et al. 2000). Intriguingly, the annotation for *Brca2* is missing in the current gerbil genome annotation, likely due to extremely unusual base composition of the genomic region in which it is found (Hargreaves et al. 2017; Pracana et al. 2020; Dai et al. 2020). While the existence and functionality of the gerbil version of *Brca2* has not yet been assessed, it is a suggestive that *Brca2* is unidentifiable in a species with a high number of rearrangements. We would not be surprised to find that the elevated mutation rate in gerbil *Brca2* underlies the elevated rate of rearrangements by impairing the function of homologous recombination and forcing gerbils to rely on non-homologous end joining. In addition, the two species with the most rearrangements (*Meriones* and *Microtus*) are unique among our focal species in having a negative relationship between recombination rate and the distance from a breakpoint (Fig. S4). This is consistent with the hypothesis that sequence variation in DNA repair pathways drive karyotypic evolution. Indeed, inter-species variability in germline double strand break repair may prove to be of greater significance to our understanding of genomic rearrangements than sequence-level genomic features. It is important to note that we propose this model as a potential mechanism that warrants further research, rather than the definitive answer to explain genome rearrangements.

In conclusion, we have found little to no support that any type of genome feature is enriched in breakpoint regions. Based on the literature, the forces that govern karyotypic evolution appear to vary drastically by lineage (Table 1, S1) and indeed we can find no consistent signal across even relatively closely related species in terms of what genome features correlate with breaks. Our results call into question the utility of looking for genome-level correlates of the location of rearrangments. We instead suggest that karyotypic evolution may be driven by sequence variation at one or a few genes that control the repair pathway used to fix double-stranded DNA breaks. Sequence variation in a few genes is a simple explanation that could explain the heterogeneity in rates of karyotypic evolution across the tree of life.

## Methods

### Genomes and genetic maps

Our focal species were three murid rodents: house mouse (*Mus musculus*), rat (*Rattus norvegicus*), and Mongolian gerbil (*Meriones unguiculatus*); and three cricetid rodents: deer mouse (*Peromyscus maniculatus*), meadow vole (*Microtus orchogaster*), and Chinese hamster (*Cricetulus griseus*). We chose these species because they are either high-quality chromosome-scale genomes (*Mus, Rattus*, and *Peromyscus*) or have striking patterns of genome evolution and high-quality (though not necessarily chromosome-scale) genomes (*Microtus, Meriones* and *Cricetulus*). House mouse, rat, gerbil, vole, and deer mouse have typical rodent karyotypes of 2N around 40. The Chinese hamster has experienced an excess of fusions which has resulted 2N=22 (Gamperl et al. 1976) while the meadow vole is reported to have many inversions (Romanenko et al. 2018; McGraw et al. 2011). Gerbils are reported to experience a high number of rearrangements, usually claimed to be Robertsonian translocations (Benazzou et al. 1982). In sum, we found six closely related species with a variety of patterns of genome evolution with which we hope to uncover general patterns in genome evolution. While this is a small set, it is similar in size to many of the studies that have found genomic correlates of rearrangments (see Table S1).

We downloaded target species genomes from NCBI: *Mus musculus* (house mouse, mm10, release 93, (Hunt et al. 2018)), *Rattus norvegicus* (rat, rn6.0, release 93, (Hunt et al. 2018)), *Meriones unguiculatus* (Mongolian gerbil, GCF_002204375.1, (Zorio et al. 2018)), *Microtus ochrogaster* (meadow vole, GCF000317375.1, (McGraw et al. 2011)), *Cricetulus griseus* (Chinese hamster, APMK01000000 (Brinkrolf et al. 2013)), and *Peromyscus maniculatus* (deer mouse, GCA_003704035.1). The house mouse, rat, deer mouse, and meadow vole genomes are chromosome-level assemblies. The Mongolian gerbil genome assembly has 68,793 scaffolds and a scaffold N50 of 374,687 bases (Zorio et al. 2018). The Chinese hamster scaffolds were sorted and assigned to chromosomes but unordered within each chromosome, with chromosomes 9 and 10 in the same pool. There are 28,764 scaffolds in this genome with an N50 of 1,245,000 bp. Only the Chinese hamster has not previously been annotated. We also downloaded genetic maps for house mouse (Cox et al. 2009), rat (FileS4 from (Littrell et al. 2018)), deer mouse (Kenney-Hunt et al. 2014; Brown et al. 2018), vole (‘Supplemental Table 1’ from (McGraw et al. 2011)), and gerbil (Brekke et al. 2019). No genetic map for Chinese hamster exists.

### Syntenic fragment and breakpoint identification

House mouse (repeat-masked version), deer mouse, Chinese hamster and meadow vole genomes were all aligned to the repeat-masked rat genome with mummer (Kurtz et al. 2004) using nucmer (flags: --mum, −c 20, and −g 1000), delta-filter (flags: −1 −i80 −1 500 −u 0 −o 100), and show-coords (flags: −r −1 −T). While it should not matter to the locations of breakpoints we identify, we chose the rat genome as it is the best quality (except for mouse which is thought to have an high number of rearrangements compared to the other species). It seems prudent to not use the most-rearranged genome as the baseline. For each pairwise comparison to rat, all alignments containing more than 5% Ns were removed to decrease noise and every chromosome pair that shared greater than 100,000 bases of aligned sequence were inspected for syntenic fragments and breakpoints. This threshold length is not a minimum rearrangement length but the minimum number of shared bases between two chromosomes in order for any rearrangements to be identified. Any syntenic fragment found between a pair of chromosomes may be shorter than this threshold, but two chromosomes must share at least 100k bases for any rearrangement to be considered in order to minimize spurious alignments. Syntenic fragments (SFs) were identified as stetches of DNA that remained colinear across all the species and breakpoint regions (BPs) were identified as the unaligned bases between two adjacent SFs. Therefore, while syntenic fragments are a function of the phylogeny and so the identity and order of the genes on a given SF in one species are the same as that in all the other species, the breakpoints regions are species-specific; the genes and sequence within a given BP is unique to each species. Inversions that occur within a single species were not analysed further, as such inversions can be caused by assembly errors and may not represent the actual genome sequence in nature. For example, the *Microtus* genome suffers from a high rate of false inversions (McGraw et al. 2011). False-positive inversion errors are difficult to identify bioinformatically and can grossly inflate estimates of the rearrangement rate. Instead, we focused on rearrangements between chromosomes.

Breakpoint identification was done iteratively: we began with the alignment between house mouse and rat and identified all syntenic fragments and breakpoint regions. Then we analysed the deer mouse-rat alignment and found new breakpoint regions. Deer mice have many breakpoints not found between house mouse and rat including all rearrangments that occured along the murid lineage before the mouse-rat split and the rearrangments that occured in the cricetid branch of the phylogeny. Each new breakpoint caused a split in what was previously a single syntenic fragment between mouse and rat. Next we did the same with the vole-rat alignment, and the hamster-rat alignment. With the addition of each new species the number of syntenic fragments increased as the previous set was re-evaluated in light of the additional breakpoints. Hamster-specific breakpoints were apparent when neighboring regions in rat aligned to different chromosomes in hamster. In addition, Mlynarski et al. (2009) used a chromosome staining approach to identify syntenic fragments between house mouse, rat, deer mouse, and Chinese hamster which we used to verify the order of the fragments found in the Chinese hamster genome and to distinguish between hamster chromosomes 9 and 10.

### Assembly of the fragmented genomes

We used our inferred SFs to improve the assembly of the *Meriones* and *Cricetulus* genomes. The *Meriones* genome was assembled form Illumina short reads and has a genetic map available. The *Cricetulus* genome was sequenced with short reads from sorted chromosomes, but no genetic map is available. We inferred the sequence of gerbil chromosomes in two steps. First, we built gerbil syntenic fragments by aligning gerbil scaffolds to the rat genome with mummer (Kurtz et al. 2004) as described above. We then used assemble_ECR_from_genomes.py (see Dataset S2) to build the scaffolds into syntenic fragments where the within-SF order, orientation, and relative spacing of each scaffold is inferred based on the rat genome. Syntenic fragments were then ordered using the linkage data in the genetic map (Table S1). We converted the locations of genes in the annotation file to the new SF coordinates using recoordinate_GFF_for_assembled_ECR_genome.py (see Dataset S2, Table S1). Gerbil chromosomes were built by ordering the SFs using a bwa alignment between the gerbil SFs and the genetic markers. In some cases, genetic markers from multiple linkage groups aligned to a single SF and in these cases we deduced the presence of a gerbil-specific breakpoint. Gerbil-specific breakpoints divided the SF into two as in the iterative refinement process described above and updated the original dataset. As the method relies on a linkage map, the physical location of these gerbil-specific breakpoints is approximate, and should be treated with caution. The site of the breakpoints were best approximations based on the location of genomic features that may correlate with rearrangements: for instances small inversions or simply long runs of Ns in the other species.

We aligned the unordered chromosome-sorted assembly of the Chinese hamster to rat using mummer and the hamster scaffolds into syntenic fragments as above. We used chromosome painting (Mlynarksi et al (2009)) to order the SFs within each chromosome. As with gerbils, when a hamster-specific breakpoint split a previously contiguous SF, we updated the original dataset.

### Genome features

We tested for enrichment (or de-enrichment) of recombination rate, GC content, gene length, gene density, coding sequence (CDS) density, and CTCF site density in the breakpoint regions in two complimentary ways. First we tested for differences between SFs and BPs using a linear mixed effects model with ‘BP’ or ‘SF’ as the fixed effect and species as the random effect. Second we used a regression of the feature density across the length of the SF. For the regression analysis we extracted a region of interest from both ends of each SF appropriate to the scale at which the effect might be found. The rationale for- and final size of-each region of interest is discussed as each hypothesis is presented and summarized in Table S2. Every base in each genome was assigned as part of a syntenic fragment or a breakpoint region. Syntenic fragments and breakpoint regions were divided into 1000bp windows and for each window we calculated the recombination rate, GC content, gene length, gene density, and CDS density. All genetic maps, except for the vole, included base positions as well as genetic positions facilitating calculation of recombination rate. To get the genomic coordinates for markers in the vole map we used bwa mem (Li and Durbin 2009) to align sequences to the genome and extracted the base positions of the start of each marker. Recombination rate was calculated by dividing the centimorgans between each marker by the number of megabases between the markers and that rate was assigned to each base between the markers. To get the rate for a window, we averaged the recombination rates of each base in the window thus accounting for marker-containing windows with two different recombination rates. The recombination data is highly skewed towards 0 with a long positive tail and so we used a log_10_ transformation. GC content was calculated for each window as the number of Gs and Cs divided by 1000. Gene length is the number of bases between the first base of the 5’UTR and the last base of the 3’UTR and was logio corrected prior to statistical analysis. To calculate gene density and CDS density, we annotated every genomic base as intergenic, 5’ UTR, CDS, intron, or 3’UTR from the appropriate GFF file (note: exons present in some transcripts but excised in others were still counted as CDS sequence). Gene density was calculated as the number of 5’ UTR, CDS, intron, and 3’UTR bases in the million bases centered on the midpoint of each window. CDS density was calculated as the number of CDS bases in the million bases centered on the midpoint of each window. A window size of 1 million was chosen for the gene- and CDS-density to assure that few windows contained 0 genes. As the windows step by 1000 bases, and enrichement or de-enrichment near a breakpoint region should be apparent. For house mouse we also calculated the density of CTCF sites, the locations of which we downloaded from BioMart using genome version GRCm38.p6 and the “Mouse Regulatory Features” option with the filter “CTCF Binding Site”. CTCF site density was calculated as the sum total number of bases in CTCF sites across the window divided by 1000.

To avoid multiple testing errors by running independent tests for every type of repetitive element we could think of, we instead analysed the sequence similarity of syntenic fragments versus breakpoint regions using the kmer-counting, alignment-free, sequence comparsion tool, KAST (https://github.com/martinjvickers/KAST) This software uses kmer spectra to characterize the signature of DNA sequences and will identify whether repetitive elements in general are common across breakpoint regions. If KAST finds a significant difference, we can then interrogate the breakpoint regions to identify the specific type of repetitive element present. However, KAST itself is agnostic to any specific repeat. Kmer spectra are vectors composed of the frequency of occurrence of all DNA oligonucleotides or “words” of size *k* in a DNA sequence (Zielezinski et al. 2017). A similarity score for two sequences can be obtained by calculating the distance between two vectors. We ran KAST for kmers of size 2, 3, 4, 5, 6, and 7 using the Chebyshev, d2 (i.e., the cosine of the angle between the frequency vectors), and Euclidian distances. Within each chromosome, we measured the kmer distances between every pair of sequences and recorded whether the pair was comprised of two breakpoint regions, two syntenic fragments, or a breakpoint region and a syntenic fragment. As comparing similarly sized DNA regions is crucial (https://github.com/martinjvickers/KAST) we extracted six evenly spaced sub-fragments of 50,000 non-N bases from each syntenic fragment and breakpoint region and compared the kmer frequency distributions between them. If a BP or SF was too short to fit six sub-fragments, we took as many as possible and those with a non-N length less than 50kb were removed from this analysis. Each distance score was transformed by taking its log_10_. We used a linear mixed effects model of the transformed distances modeled on the type of pair with species and kmer length as random effects to test whether breakpoint regions are more or less similar to each other than syntenic fragments. Genome rearrangements

We used the program MGR (Bourque and Pevzner 2002) to measure the rearrangement distance between the genomes of all six focal species. A set of 115 syntenic fragments contained at least one marker in the Mongolian gerbil genetic map and were present in all other genomes. We extracted the order of these markers in each genome using a custom python script. We ran MGR twice, once phylogenetically naïve, and once phylogenetically-informed based on Timetree’s estimate of the species relationship within Muridae and Cricetidae ((*Mus, Rattus*), *Meriones*), (*Cricetulus, Microtus*), *Peromyscus*))(Kumar et al. 2017).

### Code archive

The entire code base for this paper including all program calls and all custom scripts has been formatted to be run with a single bash file called “do_it_all_Genome_ECR_analysis.sh” and it, along with all the custom python and R scripts, can be found in Dataset S2. All intermediate data frames can be found in Dataset S3.

## Supporting information

Dataset S1 Syntenic fragment coordinates

Table S1 Literature Review Detail

## Acknowledgements

The authors would like to thank Roddy Pracana, Yichen Dai, Adam Hargraves, and Peter Holland for helpful discussions pertaining to the project. TDB would like to thank Kris Crandell.

## Statements and Declarations

This work was supported by the Leverhulme Trust grant entitled “Decoding Dark DNA” (grant number RPG-2018-433) to J.F.M and by the Natural Environmental Research Council of the UK (grant number NE/R001081/1 to A.S.T.P). The authors declare no conflicts of interest pertaining to this manuscript and the analysis presented.

## Data availability Statement

All primary data for this manuscript are publicly available from NCBI (genomes, annotations), BioMart (CTCF sites), or in association with the referenced manuscript in which it was published (genetic maps). In addition, we have uploaded our intermediate tables containing the data we extracted from the primary sources and the code we used to do so in Dryad (doi:10.5061/dryad.r2280gbd7).

Temporary Dryad link for reviewers here: https://datadryad.org/stash/share/2ksbMvf4EJ0xvsYTANA2WJS_WDI-ha8pIpB_u94jKz4.

## Author Contributions

T.D.B, J.F.M, and A.S.T.P. concieved the experiment. T.D.B did the analysis. M.T.S wrote the code to analyse the kmer frequency. T.D.B wrote the manuscript. All authors edited and reveiwed the manuscript.

